# The activation of pentose phosphate pathway flux by hydrogen peroxide is not regulated by NADPH-mediated feedback inhibition

**DOI:** 10.1101/2025.03.17.643767

**Authors:** C. Aburto, V. Parada-Goddard, A. San Martín

## Abstract

**Background and Purpose:** Oxidative stress induces a rerouting of metabolic flux from glycolysis to the pentose phosphate pathway. One proposed mechanism involves negative feedback via tonic inhibition of glucose-6-phosphate dehydrogenase by NADPH. However, recent evidence shows that NADPH levels do not decrease five seconds after hydrogen peroxide (H_2_O_2_) treatment. This finding is inconsistent with the canonical model wherein feedback inhibition loop is modulated by NADPH-depletion. This inconsistency prompts us to test the involvement of feedback inhibition at high temporal resolution.

**Experimental Approach:** We employed genetically encoded fluorescent indicators for H_2_O_2_ (HyPerRed) and NADPH (iNap1) expressed in epithelial HEK293 cells. These tools enable simultaneous real-time, single-cell monitoring of NADPH and H_2_O_2_.

**Key Results:** Glucose sustains NADPH levels under acute oxidative stress in the first seconds following H_2_O_2_ exposure. This result contradicts the reported feedback inhibition, which is considered one of the fundamental mechanisms to explain the acute rerouting of glycolysis to PPP. Furthermore, pharmacological inhibition of G6PDH suggests that the PPP is the primary source of cytosolic NADPH under oxidative stress. Monitoring NADPH levels following G6PDH inhibition allows for the assessment of the NADPH consumption flux. This parameter is low under baseline conditions, but rises dramatically under oxidative stress.

**Conclusion and Implications:** Our results support an anticipatory phenomenon that maintains NADPH levels under acute H_2_O_2_ exposure, thereby discarding the proposed feedback inhibition loop. This work offers a new perspective on the regulatory nuances of a metabolic pathway implicated in aging, cancer and a plethora of pathological conditions associated with the deleterious consequences of oxidative stress.

## INTRODUCTION

The pentose phosphate pathway (PPP) is the main source of reducing power of the cell in the form of NADPH. It consists of a series of catabolic reactions branching from glycolysis at glucose-6-phosphate and comprises two major phases: oxidative and non-oxidative. The oxidative branch includes three irreversible reactions, and its primary product is NADPH. In the first reaction, glucose-6-phosphate (G6P) is dehydrogenated by glucose-6-phosphate dehydrogenase (G6PDH) to yield NADPH and 6-phosphogluconolactone (6PGL). In the second reaction, catalyzed by 6-phosphogluconolactonase (6PGLH), 6PGL is converted to 6-phosphogluconate (6PG). In the third step, 6PG is converted to ribulose-5-phosphate (Ru5P), producing a second molecule of NADPH via 6-phosphogluconate dehydrogenase (6PGDH). Thus, the oxidative branch serves as the primary source of antioxidant capacity in the form of NADPH, which is critical for replenishing reduced glutathione (GSH) consumed during the scavenging of <u>R</u>eactive <u>O</u>xygen <u>S</u>pecies (ROS).

ROS refers to molecules derived from molecular oxygen formed by redox reactions or electronic excitation. Hydrogen Peroxide (H_2_O_2_) is the primary ROS in redox regulation of biological activities (Sies & Jones, 2020; Winterbourn, 2018) and exerts its functions by oxidizing thiolate groups from cysteines in target proteins. H_2_O_2_ concentrations range from 1 to 500 nM in health and disease (Lyublinskaya & Antunes, 2019; Sies & Jones, 2020). This wide range of concentrations underlies its pleiotropic effects, spanning from cell signaling to oxidative damage in cancer, inflammation, and cell death. Consequently, steady-state H_2_O_2_ levels are tightly controlled by both generator and removal systems. The main sources of H_2_O_2_ includes transmembrane NADPH oxidases, which account for about 45% (Bedard & Krause, 2007), and the mitochondrial electron transport chain at complex I, II within the mitochondrial matrix, and complex III within the intermembrane space, totaling around 40% (Sies & Jones, 2020; Wong, Benoit & Brand, 2019). Its removal is carried out by both enzymatic and non-enzymatic systems. Enzymatic systems such as catalase, glutathione peroxidase, thioredoxin, peroxiredoxin, and glutathione transferase. Additionally, non-enzymatic systems, including vitamins, cofactors, and glutathione (GSH). These mechanisms have evolved to act within seconds, ensuring a rapid defense against oxidative stress. Under this condition reduced glutathione (GSH) is quickly oxidized (GSSG) and subsequently restored by GSSG reduction, using NADPH as the electron donor in a reaction catalyzed by a GSH reductase. As a result, H_2_O_2_ metabolism is rapidly and tightly regulated, with NADPH serving as the principal electron donor to counteract acute oxidative stress.

It has been proposed that H_2_O_2_ induces a rerouting of metabolic flux from glycolysis to PPP. The proposed mechanisms include the inhibition of pyruvate kinase M2 (PKM2) in cancer cells (Anastasiou et al., 2011) or glyceraldehyde 3-phosphate dehydrogenase (GAPDH) in baker’s yeast (Ralser et al., 2007; Ralser, Wamelink, Latkolik, Jansen, Lehrach & Jakobs, 2009) and mammalian cells (Colussi, Albertini, Coppola, Rovidati, Galli & Ghibelli, 2000) through the oxidation of key cysteine residues. This process leads to passive accumulation of PPP intermediates, which promotes increased carbon flux through the PPP and thereby enhances NADPH production. Alternative evidence suggests that the PPP is directly regulated by oxidative stress. This is supported by results from *in vitro* biochemical experiments, showing that NADPH inhibits purified human erythrocyte G6PDH with a Ki of 16 µM (Luzzatto, 1967). This finding is relevant because the reported cytosolic NADPH concentration in mammalian cells is close to the Ki of G6PDH, 3.1 µM (Tao et al., 2017; Zou et al., 2018). Thus, increased NADPH consumption resulted in a decrease in its concentration, relieving the inhibition and unleashing G6PDH activity. This suggests a NADPH-mediated feedback inhibition, that may represent a physiological mechanism for regulating the PPP (Eggleston & Krebs, 1974; Holten, Procsal & Chang, 1976; Luzzatto, 1967). Currently, two alternative mechanisms based on the allosteric inhibition of G6PDH by NADPH have been proposed, to explain the acute shift of carbon flux from glycolysis into the PPP. In the first mechanism, H_2_O_2_ causes a reduction in NADPH levels, which relieves inhibition on G6PDH and increases flux through the PPP, as evidenced by elevated 6-phosphogluconate (6PG) (Christodoulou, Link, Fuhrer, Kochanowski, Gerosa & Sauer, 2018). The accumulation of 6PG further amplifies this rerouting by inhibiting glucose-6-phosphate isomerase (PGI) activity (Kuehne et al., 2015). Such conclusions were drawn using time-resolved metabolomics and ultra-short ^13^C labeling experiments that allow sample acquisition times as fast as minutes. Alternatively, NADPH depletion unleashes G6PDH activity by releasing its tonic inhibition. This phenomenon was first described in bacteria (Christodoulou, Link, Fuhrer, Kochanowski, Gerosa & Sauer, 2018) but is also present across different organisms (Christodoulou, Kuehne, Estermann, Fuhrer, Lang & Sauer, 2019). H_2_O_2_-induced NADPH depletion has been demonstrated via HPLC-mass spectrometry coupled with dynamic isotope tracing with five seconds resolution, alongside computational modeling to corroborate NADPH’s role in regulating PPP flux (Christodoulou, Link, Fuhrer, Kochanowski, Gerosa & Sauer, 2018). Both proposed mechanisms are independent and precede the H_2_O_2_-driven inhibition of GAPDH or PKM2. Therefore, NADPH is crucial due to its allosterically control of PPP flux on a timescale of seconds to minutes through a feedback inhibition loop.

NADPH-mediated feedback inhibition mechanism requires a NADPH depletion. However, recent data on NADPH dynamics, measured by metabolomics and imaging approaches, show either increase or unchanged NADPH levels five seconds or minutes after exposure to 80 µM, 1 mM or sub-lethal levels of H_2_O_2_ (Christodoulou, Link, Fuhrer, Kochanowski, Gerosa & Sauer, 2018; Nikel, Fuhrer, Chavarria, Sanchez-Pascuala, Sauer & de Lorenzo, 2021; Tao et al., 2017).

These results contradict the previously proposed NADPH-dependent feedback inhibition mechanism, which needs a fast decrease of NADPH to initiate the regulatory loop. In this work, we have tested the presence of a feedback inhibitory loop at high temporal resolution and propose an alternative explanation for PPP regulation by H_2_O_2_, mediated by a feedforward phenomenon that increases NADPH production before its depletion. To test this, we leveraged the minimally invasive, multiplexing capabilities and high spatiotemporal resolution of Genetically Encoded Fluorescent Indicators for NADPH (iNap1) and H_2_O_2_ (HyPerRed).

## METHODS

### Cell Culture and Transfection

HEK293 cells were acquired from the American Type Culture Collection (ATCC) and cultured at 37°C in 95% air/5% CO_2_ in DMEM/F12 10% fetal bovine serum. Cells were co-transfected with 1 µg iNap1/pcDNA3.1 and HyPerRed/pC1 plasmid DNA of at 60% confluency using Lipofectamine 3000 (Gibco). This was incubated for 16 hours with an efficiency of >60% and imaged at room temperature (22−25 °C).

### Fluorescence measurements

For cell imaging experiments, cells were superfused with a solution of the following composition (in mM): 136 NaCl, 3 KCl, 1.25 CaCl_2_, 1.25 MgSO_4_, 10 HEPES pH 7.4, using an upright Olympus FV1000 confocal microscope equipped with a 20x water immersion objective (N.A. 1.0). 440 and 543 nm solid-state lasers were used to image iNap1 and HyPerRed, respectively. Time series images were taken every 1 s with 20× (NA 1.0) in XYT scan mode (scan speed: 125 kHz; 128 × 128-pixel; pinhole 800 μm).

### Mathematical modelling of feedback

The predicted response of the feedback between G6PDH and NADPH was simulated using Berkeley Madonna software and the following set of ordinary differential equations:

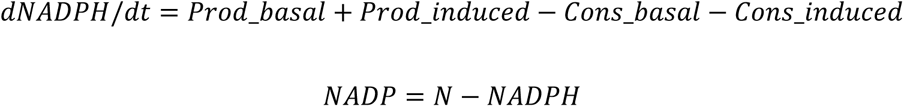

where Prod_basal is the production through G6PDH in the absence of peroxide, Prod_induced is the increase in production flux in response to feedback, Cons_basal is the NADPH consumption in the absence of H_2_O_2_, Cons_induced is the increase in NADPH consumption flux in response to the addition of H_2_O_2_, and N is the total concentration of NADPH and NADP^+^. N was fixed at 3 µM (Tao et al., 2017), Prod_basal was set to 0.083 µM/s, and Cons_basal was defined as k_cons * NADPH, where k_cons was adjusted to 0.0147. Both values were calibrated to obtain an NADPH concentration of 2.97 µM, considering a NADPH/NADP^+^ ratio of 100 at rest (Merrill & Guynn, 1981). Cons_induced was defined as a constant flux of 3 µM/s, independent of basal consumption, activated as a square function simulating the acute presence of H_2_O_2_.

Prod_induced was defined as (V_max_ * NADP^H^) / (K_m__NADP^H^ + NADP^H^) to achieve feedback sensitive to NADP^+^. To stimulate an ultrasensitive feedback response, the Hill coefficient (H) and K_m__NADP were defined with the following values: (1.69, 0.368522 µM; 2, 0.249813 µM; 4, 0.0865701 µM; 20, 0.0370826 µM). The simulations were expressed directly as NADPH concentration, so modeling of GEFIs was not necessary. The lowest Hill coefficient used in the simulations (H = 1.69) corresponds to the value described in the literature for G6PDH-NADP^+^ (Luzzatto, 1967).

### Data and statistical analysis

For data analysis, the fluorescent signal from a region of interest (ROI) from each cell was collected. Background subtraction was performed separately for each channel. Regression and statistical analysis were carried out using SigmaPlot (Jandel).

## MATERIALS

NaCl, KCl, CaCl_2_, MgSO_4_ and HEPES were obtained from Sigma (St Louis, MO, USA). H_2_O_2_ (CAS N°107209) was purchased from Merckmilipore and a stock solution (1 M) prepared in water. Sodium L-Lactate (CAS N° 867-56-1) and Sodium Pyruvate (CAS N° 113-24-6) were purchased from Sigma-Aldrich and a stock solution (1 M) prepared in water stored at 4°C. G6PDi-1 (CAS N°2457232-14-1) was purchased from Sigma-Aldrich and prepared in DMSO stock (7 mM). Diamide (CAS N°10465-78-8) was purchased from Sigma-Aldrich and a stock solution (120 mM) prepared in water.

## RESULTS

### Hydrogen peroxide acutely modulates NADPH levels

Hydrogen peroxide induces intracellular NADPH depletion in mammalian cells when carbon sources are absent after minutes of oxidative stress exposure (Tao et al., 2017). To determine the H_2_O_2_ concentration that reproducibly affects cytosolic NADPH relative levels, we performed doses-response experiments in HEK293 cells simultaneously expressing iNap1 to monitor NADPH (Tao et al., 2017) and HyPerRed to monitor H_2_O_2_ (Ermakova et al., 2014) (**Figure 1A**). Increasing concentrations of exogenous H_2_O_2_ produced a rapid decline in NADPH levels, with a robust and reproducible increase of intracellular H_2_O_2_ to a 250 µM pulse in carbon-source-starved conditions (**Figure 1B,C**). At this extracellular concentration of H_2_O_2_, the effective intracellular concentration is approximately 400 nM, based on reported gradients between extracellular and intracellular milieu (Huang & Sikes, 2014; Lyublinskaya & Antunes, 2019).

**FIGURE 1.**
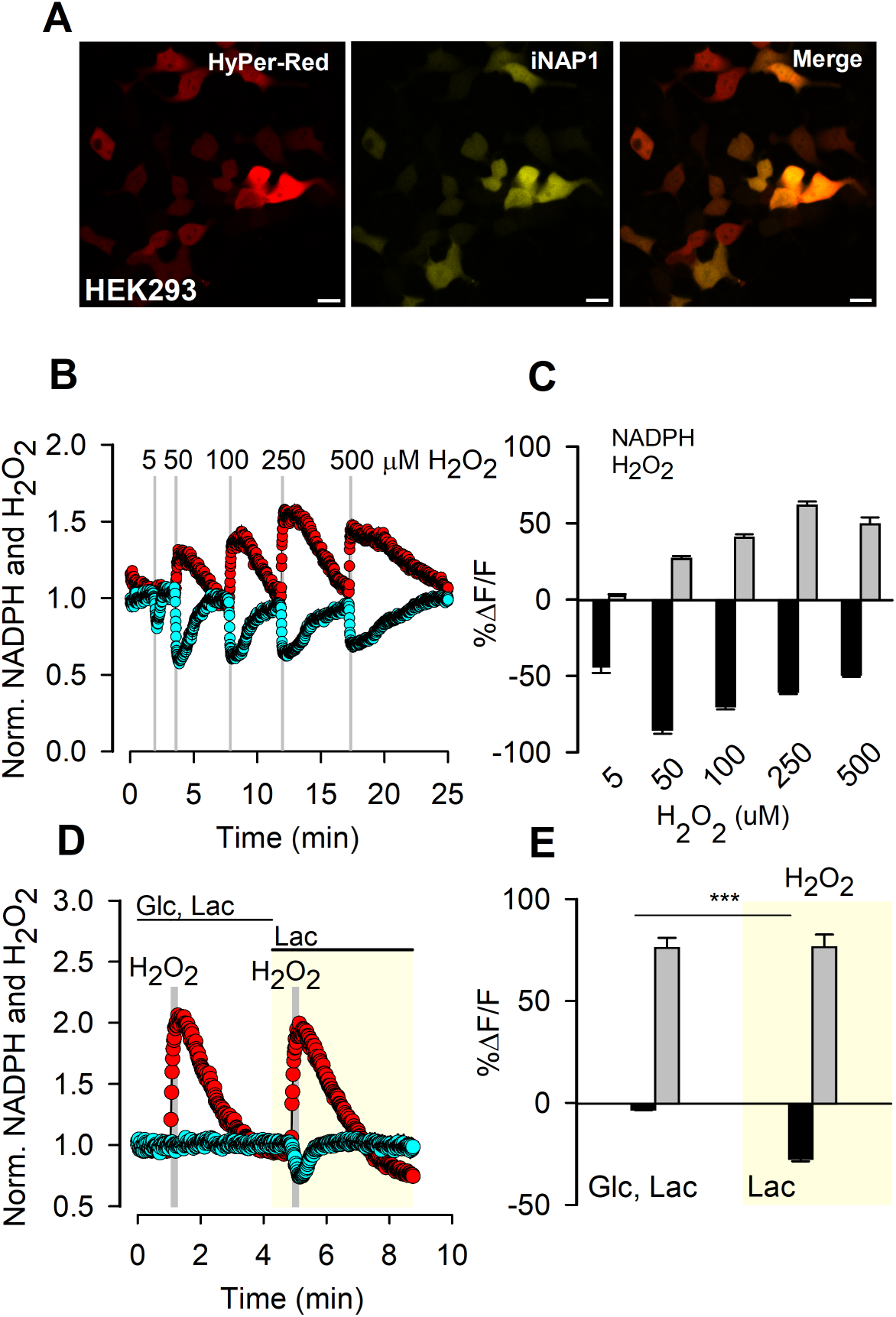
Hydrogen peroxide acutely modulate NADPH levels. **(A)** Representative images of HEK293 cells co-expressing HyPerRed (red) and iNap1 (yellow), along with their merged signals. Scale bar: 20 µm. **(B)** Representative normalized fluorescence traces of iNap1 (cyan circles) and HyPerRed (red circles) in response to 5, 50, 100, 250, and 500 µM H_2_O_2_ during ∼30 seconds each. Acquisition interval was 1 second. **(C)** Summary bars of fluorescence changes in NADPH (black) and H_2_O_2_ (gray) levels in response to increasing H_2_O_2_ concentrations. Data is presented as mean ± SEM from three independent experiments (n = 30 cells). **(D)** Representative normalized fluorescence traces of iNap1 and HyPerRed in response to two pulses of 250 µM H_2_O_2_, in the presence of either 5 mM glucose and 1 mM lactate or only 1 mM lactate (highlighted in light yellow). Acquisition interval was 1 second. **(E)** Summary bars of fluorescence changes in NADPH (black) and H_2_O_2_ (gray) levels following treatment with 250 µM H_2_O_2_ in presence of 5 mM glucose and 1 mM lactate or only 1 mM lactate. Data is presented as mean ± SEM from three independent experiments (n = 33 cells, ***p < 0.001, two-tailed paired t-test).

Previous results demonstrated that in glucose-feed cells, NADPH levels remain unperturbed (Scherschel et al., 2024; Tao et al., 2017) because of increased flux through the PPP (Christodoulou, Link, Fuhrer, Kochanowski, Gerosa & Sauer, 2018), thereby enhancing NADPH production. However, these experiments were performed at low temporal resolution, which precluded observation of the acute effect of H_2_O_2_. To investigate whether glucose is critical to sustain NADPH levels on a timescale of seconds, we conducted experiments in the presence and absence of glucose while simultaneously monitoring NADPH and H_2_O_2_ (**Figure 1D**). A short pulse of H_2_O_2_ in the presence of carbon sources such as glucose and lactate did not induce a NADPH depletion (**Figure 1D,E**). Moreover, when we removed the glucose from the extracellular buffer, lactate was unable to maintain steady-state NADPH levels under acute H_2_O_2_ stress (**Figure 1D,E**). These results confirm that NADPH levels remain unperturbed under acute H_2_O_2_ stress when glucose is provided as a carbon source, and highlight glucose as a necessary carbon source to sustain the NADPH steady-state under oxidative stress.

### Hydrogen peroxide does not cause an immediate change in NADPH levels

The proposed NADPH-mediated feedback inhibition mechanism implies a significant decrease in NADPH levels. However, our previous results and recent data on NADPH dynamics that were measured by metabolomics and imaging approaches, showed no reduction in NADPH levels after H_2_O_2_ treatment in the presence of carbon sources. To determine whether H_2_O_2_ induces an immediate NADPH depletion, we monitored NADPH and H_2_O_2_ at a second-by-second resolution in presence and absence of carbon sources. HEK293 cells expressing iNap1 and HyPerRed were exposed to a 250 µM H_2_O_2_ pulse in presence of glucose and lactate. Under these conditions H_2_O_2_ did not induce NADPH depletion during the first five seconds after its exposure (**Figure 2A,B**). In the absence of any carbon source, H_2_O_2_ induces NADPH depletion detected at 5 seconds post-exposure (**Figure 2C,D**). These results indicate that H_2_O_2_ is incapable of causing immediate NADPH depletion in the presence of glucose, ruling out NADPH as the primary signal in the reported NADPH-mediated feedback inhibition mechanism.

**FIGURE 2.**
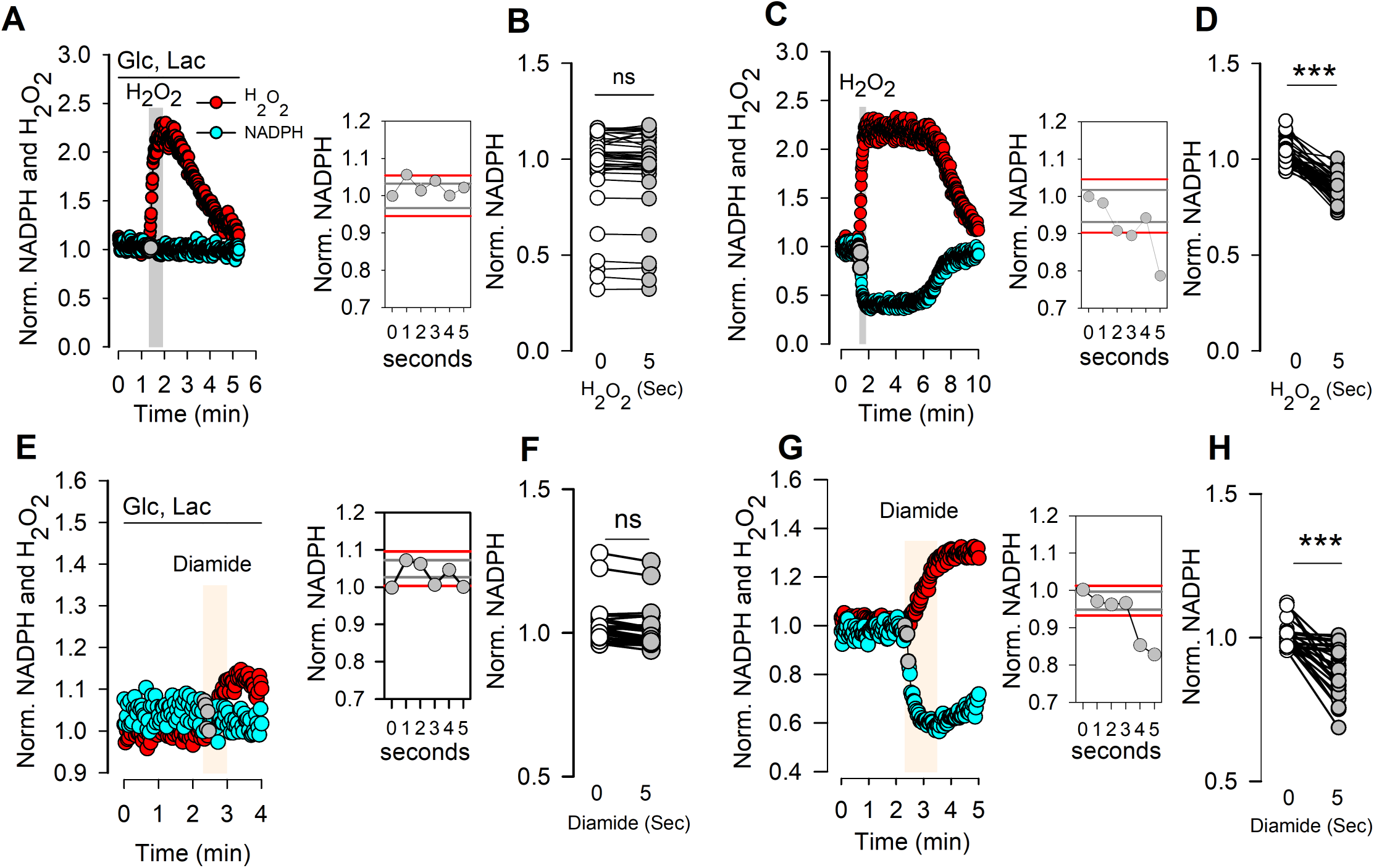
Hydrogen peroxide does not cause an immediate change in NADPH levels. **(A, C)** Representative single-cell recordings of iNap1 (cyan circles) and HyPerRed (red circles) in response to a 250 μM H_2_O_2_ pulse (∼30 s), in the presence **(A-B)** or absence **(C-D)** of 5 mM glucose and 1 mM lactate. **(E, G)** Representative single-cell recordings of iNap1 (cyan circles) and HyPerRed (red circles) in response to a 50 μM diamide pulse (∼30 s), in the presence **(E-F)** or absence **(G-H)** of 5 mM glucose and 1 mM lactate. The first 5 seconds after each perturbation are highlighted with gray circles. The insets show these first five seconds after each perturbation (gray circles). Red lines represent the 99% confidence interval, and gray lines represent the 95% confidence interval. Data was acquired every second. **(B, D, F, H)** Single-cell comparison of normalized NADPH levels at 0 seconds and 5 seconds post H_2_O_2_ **(B, D)** or diamide **(F, H)**, under the presence **(B,F)** or absence **(D, H)** of carbon sources. Data in (B) are from four independent experiments (n = 40 cells), and data in (D) are from five independent experiments (n = 50 cells). A paired t-test was performed. ns, not significant; ***p < 0.001.

To distinguish between a general oxidative stress response and unrelated effects such as those derived from H_2_O_2_ metabolism, we used diamide as a parallel oxidative challenge (Kosower, Kosower, Wertheim & Correa, 1969). Diamide selectively oxidizes thiol groups (–SH) to disulfides (–S–S–) in proteins, thereby serving as a control to confirm that the observed changes can be attributed specifically to H_2_O_2_-mediated protein oxidation. HEK293 cells expressing iNap1 and HyPerRed were exposed to a 50 µM diamide pulse in the presence of glucose and lactate monitoring at a second-by-second resolution. Under these conditions, diamide did not induce NADPH depletion during the first five seconds (**Figure 2E,F**). In the absence of any carbon source diamide was able to induce an acute decrease in NADPH levels at four seconds post exposure (**Figure 2G,H**). Therefore, these results from both H2O2 and diamide in presence of glucose and lactate rule out NADPH as the signal involved in the reported NADPH-mediated feedback inhibition, and instead provide evidence that an alternative phenomenon mediated by thiol protein modifications is involved.

### Hydrogen peroxide induces an acute increase in NADPH consumption flux

H_2_O_2_ did not induce an acute perturbation of the steady-state levels of NADPH in presence of carbon sources, a finding that conflicts with the proposed NADPH-mediated feedback inhibition. An alternative explanation is an anticipatory phenomenon in which NADPH production increases to match its consumption. This tight coupling between NADPH production and consumption would need to be highly synchronized to explain the absence of any perturbation in NADPH steady-state during the first seconds of H_2_O_2_-induced oxidative stress.

Steady-state cytosolic NADPH levels result from the net balance of its production and consumption fluxes. Main cytosolic sources of NADPH include the PPP, the malic enzyme 1 (ME1) that catalyzed conversion of malate to pyruvate, and cytosolic isocitrate dehydrogenase (IDH1) that catalyzed the conversion of isocitrate to alpha-ketoglutarate. Meanwhile, NADPH is consumed in various catabolic and anabolic reactions and in the regeneration of oxidized glutathione generated during oxidative stress. Consequently, pharmacological blockage of a single production or consumption pathway does not allow straightforward dissection of individual fluxes at single-cell resolution using iNap1, because NADPH production and consumption reactions are multiple. Obtaining such flux information is crucial, given that current methods for measuring PPP flux lack high spatiotemporal resolution.

Surprisingly, by using a previously described G6PDH inhibitor, we were able to dissect NADPH consumption flux at single cell level. HEK293 cells expressing iNap1 and HyPerRed were exposed to 7 µM of G6PDi-1, a known G6PDH with an IC_50_ of 0.07 µM (Ghergurovich et al., 2020). In the presence of carbon sources, H_2_O_2_ did not induce NADPH depletion; however, when the G6PDH inhibitor was added to block the entry into the PPP, H_2_O_2_ triggered a rapid drop in NADPH levels (**Figure 3A,B**). Based on these results, we hypothesize that oxidative stress, under the pharmacological inhibition of G6PDH, will decrease NADPH levels even in the presence of carbon sources. This outcome suggests that the alternative cytosolic sources of NADPH-producing pathways cannot compensate for the increase of its consumption (**Figure 3C,D**). Notably, a comparison of NADPH depletion rates under absence of carbon sources showed no significant difference from the depletion rate observed with the G6PDH inhibitor in presence of carbon sources (**Figure 3C,D**). This data suggests that the PPP is the main source of NADPH. Therefore, a plausible explanation for maintaining NADPH steady-state during the first seconds after acute oxidative stress is an increment of its production.

**FIGURE 3.**
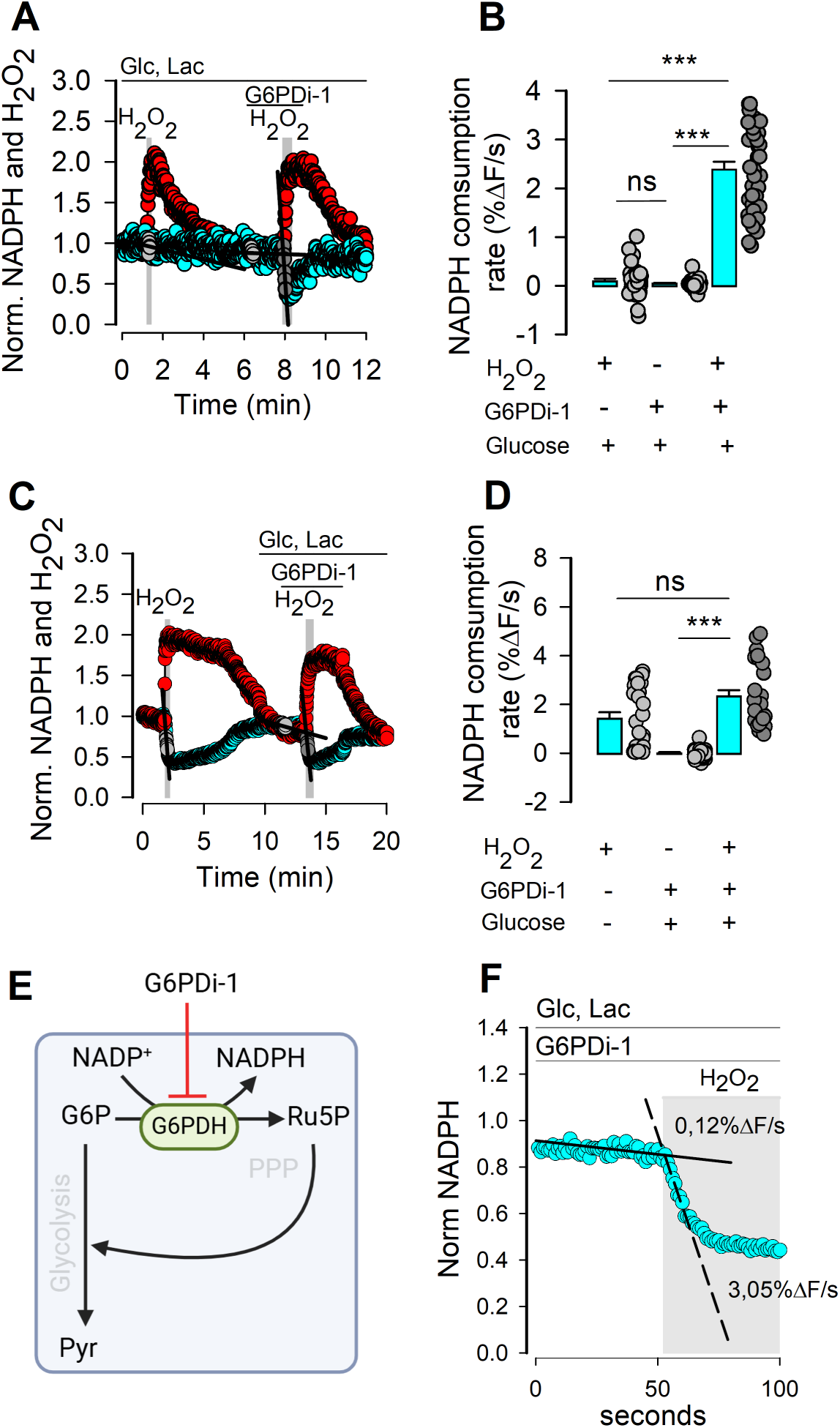
Hydrogen peroxide induced an acute increase in NADPH consumption flux. **(A)** Representative single-cell recordings of iNap1 (cyan circles) and HyPerRed (red circles). Two pulses of 250 µM H_2_O_2_ were applied in the absence and presence of 7 µM G6PDi-1, under 5 mM glucose and 1 mM lactate. Black lines and gray circles represent the traces used to calculate consumption rates. Data was acquired every second. **(B)** Comparison of NADPH consumption rates with and without H_2_O_2_ in the presence or absence of 7 µM G6PDi-1, under carbon-source conditions (5 mM glucose and 1 mM lactate). Bars represent the mean ± SEM, and gray dots show the consumption rate for each cell (n = 30 cells, 3 independent experiments). Kruskal-Wallis tests were performed, followed by Dunn’s post hoc test. ns, not significant; ***: p < 0.001. **(C)** Representative single-cell recordings of iNap1 (cyan circles) and HyPerRed (red circles). A 250 µM H_2_O_2_ pulse was applied in the absence of glucose. In the presence of glucose, 7 µM G6PDi-1 was added, followed by another 250 µM H_2_O_2_ pulse. Black lines and gray circles represent the traces used to calculate consumption rates. **(D)** Comparison of NADPH consumption rates with and without H_2_O_2_, either in the absence of carbon sources or in the presence of carbon sources plus G6PDi-1. Bars represent the mean ± SEM, and gray dots show the consumption rate for each cell (n = 30 cells, 3 independent experiments). Friedman tests were performed, followed by Dunn’s post hoc test. ns, not significant; ***p < 0.001. **(E)** Schematic representation of NADPH production inhibition by G6PDi-1. G6PDH: Glucose-6-phosphate-dehydrogenase; G6P: Glucose-6-Phosphate; Ru5P: Ribulose-5-Phosphate; Pyr: pyruvate. **(F)** Representative inset of normalized iNap1 data presented in (A). Shows the G6PDi-1–mediated NADPH consumption rates before (solid line) and after (dash line) H_2_O_2_ addition under presence of carbon sources.

The pharmacological inhibition of G6PDH can be used to measure basal and H_2_O_2_-activated NADPH consumption (**Figure 3E**). Consistently, under basal conditions without oxidative stress, the G6PDH inhibitor does not cause NADPH depletion, indicating a low NADPH consumption rate. However, this rate accelerates significantly (from 0.12 to 3.05 %ΔF/seg) upon the exposure to H_2_O_2_ (**Figure 3F**), implying that PPP-derived NADPH is crucial for acute protective response to oxidative stress.

### Negative-feedback inhibition does not explain NADPH dynamics under H₂O₂ stress

To corroborate the feasibility of feedback inhibition as the main mechanism responsible for maintaining invariant NADPH levels following acute oxidative stress with H_2_O_2_, we constructed a numerical model simulating feedback inhibition of NADPH production in response to an invariant concentration of G6P (**Figure 4A**).

**FIGURE 4.**
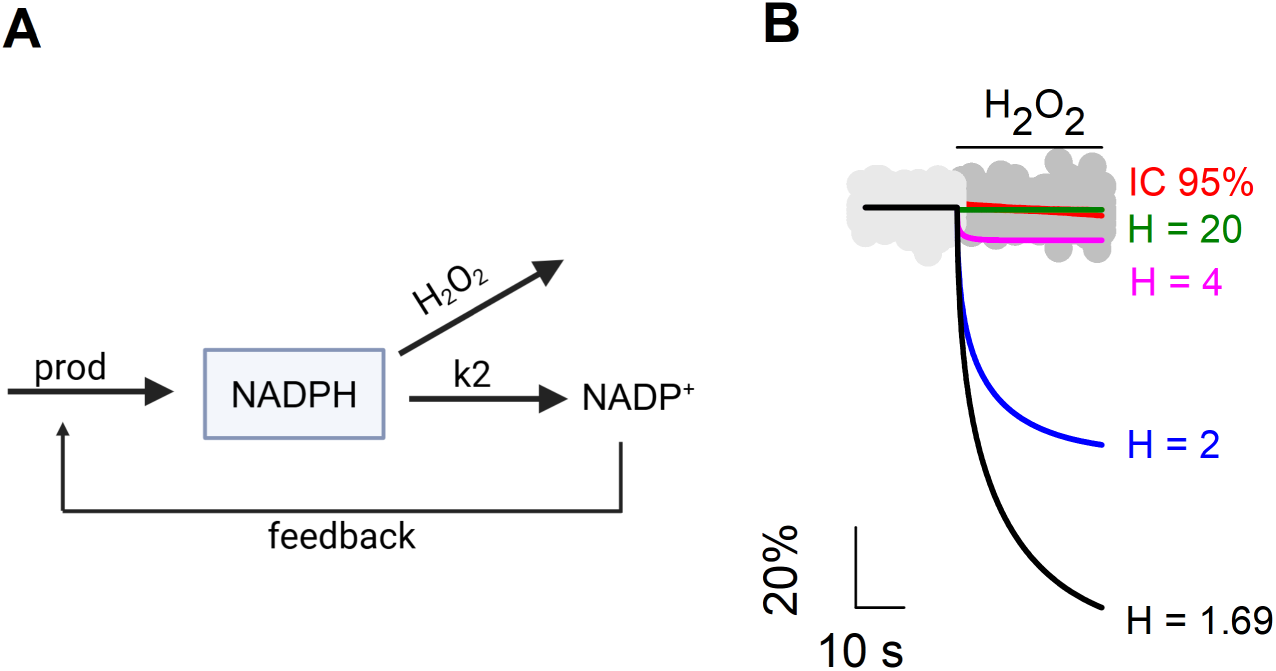
Negative-feedback inhibition does not explain NADPH dynamics under H_2_O_2_. **(A)** Schematic representation of the model used to simulate negative feedback inhibition of NADPH production. In this model, “prod” represents a constant basal flux of NADPH production, “k_2_” corresponds to the rate constant for basal NADPH consumption, “H_2_O_2_” represents NADPH consumption activated by H_2_O_2_, and “feedback” refers to the activation of NADPH production dependent on NADP⁺. **(B)** Simulation of negative feedback regulation of NADPH production in response to oxidative stress induced by H_2_O_2_, using Hill coefficients (H) of 1.69 (black), 2 (blue), 4 (magenta), and 20 (green). Data is expressed as the percentage change in NADPH concentration based on measurements with iNap1. Solid lines represent simulated data, while circles correspond to experimental data from 10 cells before (light gray) and after (dark gray) a 250 µM H_2_O_2_ pulse in the presence of 5 mM glucose. The 95% confidence interval (CI) for the mean is shown in red.

NADPH production was modeled as a basal production flux dependent on G6P. This flux was kept constant to isolate the effect of feedback inhibition. Additionally, a production flux dependent on NADP^+^ was included to represent the feedback of NADPH on G6PDH activity. This flux was defined using a Michaelis-Menten equation with a variable Hill coefficient (H) for NADP^+^, indicating different levels of G6PDH sensitivity to NADP^+^ concentration. Thus, a decrease in NADPH, which naturally leads to an increase in NADP^+^ levels, would activate this flux, as described in the feedback inhibition model.

NADPH consumption was expressed as two independent fluxes: one corresponding to basal consumption, modeled as a linear function dependent on NADPH, and an acute consumption flux induced by H_2_O_2_, defined as a fixed consumption flux that is activated in the presence of H_2_O_2_ (**Figure 4A**). The basal production and consumption values were adjusted to achieve a steady-state NADPH concentration of 3 µM 28581494 and an NADPH/NADP+ ratio of 100 (Merrill & Guynn, 1981; Veech, Eggleston & Krebs, 1969) in the absence of H_2_O_2_. Several system configurations were simulated, varying the value of H and the corresponding change in the Km of NADP^+^ to ensure a similar steady state in all cases. H values ranged from 1.69, corresponding to the H reported for G6PDH-NADP^+^ (Luzzatto, 1967), to higher coefficients, increasing until reaching an H value that allowed the curve to remain within the experimentally observed NADPH steady state in the presence of glucose after adding H_2_O_2_ (95% confidence interval) (**Figure 4B**).

The results showed that the minimum H value required to prevent a significant decrease in NADPH after a 250 µM H_2_O_2_ pulse was H = 20 (**Figure 4B**). Typical H values range between 1 and 4 (Prinz, 2010), with some cases reaching 10 and a few exceptions nearing 15 (Bai et al., 2010; Cluzel, Surette & Leibler, 2000). However, despite synthetically achieving very high H values, an H = 20 implies a sensitivity that is practically impossible to attain in physiological systems. Furthermore, since the H value for this system has been previously measured and reported as H = 1.69 (Luzzatto, 1967), it is highly unlikely that a feedback inhibition mechanism regulating NADPH production is the primary factor maintaining a constant NADPH steady state after acute oxidative stress.

## DISCUSSION

Previous reports proposed a NADPH-based feedback inhibition for the rapid control of PPP flux under oxidative stress. This mechanism describes acute response by releasing the tonic inhibition of G6PDH by NADPH, which is rapidly consumed under oxidative stress. However, current experimental data does not support this type of regulation, because a prompt drop in NADPH levels would be necessary to align with such a feedback mechanism. Instead, our second-by-second measurements reveal that H_2_O_2_ does not cause an immediate decrease in NADPH levels, thereby ruling out this cofactor as the signal that triggers the proposed feedback inhibition.

The absence of any acute perturbation in the steady-state of NADPH level by oxidative stress suggests a close coupling between NADPH production and consumption. Therefore, any increase in its consumption triggers a corresponding increase in its production. However, based on experiments performed using high H_2_O_2_ concentrations, even this rapid increase in NADPH production can become insufficient to meet the high demand for reducing power, ultimately causing NADPH depletion within a time frame of minutes to hours. Therefore, we cannot rule out the possibility of feedback inhibition occurring under conditions of overwhelming oxidative stress, sufficient to induce the observed NADPH depletion in long timeframes (Kuehne et al., 2015; Ralser et al., 2007). It is important to note that in our experiments, oxidative stress was induced by a thirty second pulse of H_2_O_2_ exposure, simulating acute treatment. Additionally, we analyzed our data before reaching the maximal H_2_O_2_ concentration generated by the pulse, meaning that we observed the expected phenomenon at lower H_2_O_2_ concentrations. This approach differs from previous studies, where H_2_O_2_ exposure typically lasted from minutes to hours and reached maximal intracellular concentration of H_2_O_2_ (Kuehne et al., 2015; Nikel, Fuhrer, Chavarria, Sanchez-Pascuala, Sauer & de Lorenzo, 2021; Ralser et al., 2007).

To maintain unperturbed NADPH steady-state levels, an anticipatory response is required to boost NADPH production before its depletion. Our data confirm that glucose is necessary for sustaining NADPH levels under oxidative stress. Although these results are consistent with previous work (Christodoulou, Kuehne, Estermann, Fuhrer, Lang & Sauer, 2019; Tao et al., 2017), they also suggest that, to meet the high demand for reducing power, oxidative agents such as H_2_O_2_ may modulate glucose transport to ensure the carbon flux necessary for the PPP and NADPH production. In this regard, it is important to investigate, on the timescale of seconds, the coupling between glucose metabolism and NADPH dynamics. Indeed, some reports have shown that H_2_O_2_ can induce an increase of glucose transport after prolonged H_2_O_2_ exposure (Hamrahian, Zhang, Elkhairi, Prasad & Ismail-Beigi, 1999; Prasad & Ismail-Beigi, 1999). Therefore, based on this evidence we cannot rule out that the sensor of oxidative stress, that permits the anticipatory phenomenon, is located within glucose metabolism, which would allow the system to enhance glucose transport and metabolism in order to preempt NADPH depletion.

Additionally, our results indicate that, under oxidative stress, most of the glucose-derived carbon flux is directed to the PPP. Specifically, inhibiting G6PDH in presence of glucose did not cause a decline in NADPH levels under resting conditions; however, upon oxidative stress activation, a rapid NADPH depletion was observed. This rate was not significantly different from the rate obtained from resting conditions without carbon sources. These data suggest that alternative NADPH sources cannot fully compensate for NADPH demand when G6PDH is inhibited in a similar way to carbon sources-deprived conditions. These results are relevant because the fraction of carbon flux partitioned between glycolysis and the PPP remains controversial (Rodriguez-Rodriguez, Fernandez & Bolanos, 2013). Furthermore, our experiments with the pharmacological G6PDH inhibition present an opportunity to develop a protocol for dissecting the NADPH consumption flux, which can be used as a proxy for PPP flux. Current methods for assessing metabolic fluxes often rely on isotopic labeling techniques coupled with mass spectrometry, as useful as they are, have certain limitations in terms of temporal resolution and single-cell measurements capabilities. We envision that our G6PDH inhibition experiments will form the basis for a method that enables the assessment of PPP flux on a timescale of seconds and at single-cell resolution, both in resting and activated states.

To better understand the non-NADPH-depletion by oxidative stress, we built a simulation of the NADPH compensation system in response to H_2_O_2_, based on a classical feedback inhibition model (Figure 4). Our simulation suggests that in order to explain the experimental data with a feedback inhibition model, the cooperativity between G6PDH and NADPH needs to be extremely high, with a Hill coefficient (H) of at least 20. This value contrasts significantly with estimates from *in vitro* experiments, where a Hill coefficient of 1.69 has been reported (Luzzatto, 1967). Moreover, in physiological contexts, typical H values range up to 3 or 4, meaning that *H* = 20 falls outside the biologically relevant range (Cluzel, Surette & Leibler, 2000; Luzzatto, 1967; Prinz, 2010). Therefore, feedback inhibition alone is not sufficient to explain the acute compensation of increased NADPH consumption following oxidative stress. Instead, an anticipatory feedback phenomenon better fits the observed results. This type of approach has also been used to assess ATP level regulation following the rapid activation of the Na⁺/K⁺ pump (Baeza-Lehnert et al., 2019) and the acute activation of glucose transport in response to potassium (Fernandez-Moncada et al., 2021). This suggests that anticipatory feedback is a mechanism employed by the cell to rapidly respond to events that could cause undesired imbalances in critical metabolites for cellular homeostasis, such as glucose, ATP, and NADPH.

Certainly, our study presents some limitations. All our experiments were performed using supraphysiological of exogenous H_2_O_2_ concentrations. However, these concentrations are lower than those reported in most studies that use over 500 µM and the exposition time is shorter, just thirty seconds. Future experiments exploring lower and more physiological concentrations of endogenous H_2_O_2_ will be important to determine whether this anticipatory phenomenon also occurs under more physiological conditions. Additionally, this study primarily describes anticipatory behavior; hence, the underlying mechanisms remain unknown.

In summary, this work provides high temporal resolution evidence that under acute oxidative stress, cytosolic NADPH levels remain constant. This observation is not fully explained by the current model of feedback inhibition that reroutes glycolysis to the PPP. Instead, we propose an anticipatory phenomenon that under oxidative stress increases the NADPH production before its depletion.

## ACKNOWLEDGMENT

Associated grants, Beca Doctorado Nacional ANID 21190573 and Universidad San Sebastián Internal Grant USS-FIN-23-FAPE-02.

